# Genome-wide CRISPR screens reveal APR-246 (Eprenetapopt) triggers ferroptosis and inhibits iron-sulfur cluster biogenesis

**DOI:** 10.1101/2020.11.29.398867

**Authors:** Kenji M. Fujihara, Bonnie Zhang, Thomas D. Jackson, Brunda Nijiagel, Ching-Seng Ang, Iva Nikolic, Viv Sutton, Joe Trapani, Kaylene J. Simpson, Diana Stojanovski, Silke Leimuehler, Sue Haupt, Wayne A. Phillips, Nicholas J. Clemons

## Abstract

The mechanisms by which cells respond and adapt to oxidative stress are largely unknown but are key to developing a rationale for cancer therapies that target antioxidant pathways. APR-246 is a mutant-p53 targeted therapeutic currently under clinical investigation in myeloid dysplastic syndrome (MDS) and acute myeloid leukemia^1^. Whilst the mechanism of action of APR-246 is thought to be reactivation of wild-type p53 activity through covalent modification of cysteine residues in the core domain of mutant-p53 protein^2,3^, here we report that the anti-neoplastic capacity of APR-246 lies predominantly in the conjugation of free cysteine. Genome-wide CRISPR perturbation screening, metabolite profiling and proteomics in response to APR-246 treatment in mutant-p53 cancer cells highlighted the role of GSH and mitochondrial metabolism in determining APR-246 efficacy. APR-246 sensitivity was increased through loss of key enzymes in mitochondrial one-carbon metabolism, *SHMT2* and *MTHFD1L*, due to diminished glycine supply for *de novo* GSH synthesis. Critically, we show that APR-246 induces iron-dependent, apoptotic machinery-independent cell death, ferroptosis. Whole-cell proteomics analyses indicated an upregulation of proteins involved in iron-sulfur cluster biogenesis (eg. FDX1). GSH, acetyl-CoA and NADH levels were also depleted in APR-246 treated cells. Importantly, we found that APR-246 inhibits iron-sulfur cluster biogenesis in the mitochondria of cancer cells through cysteine conjugation. This work not only details novel determinants of APR-246 activity in cancer cells, but also provides a clinical roadmap for targeting antioxidant pathways in tumours - beyond targeting mutant-p53 tumours.

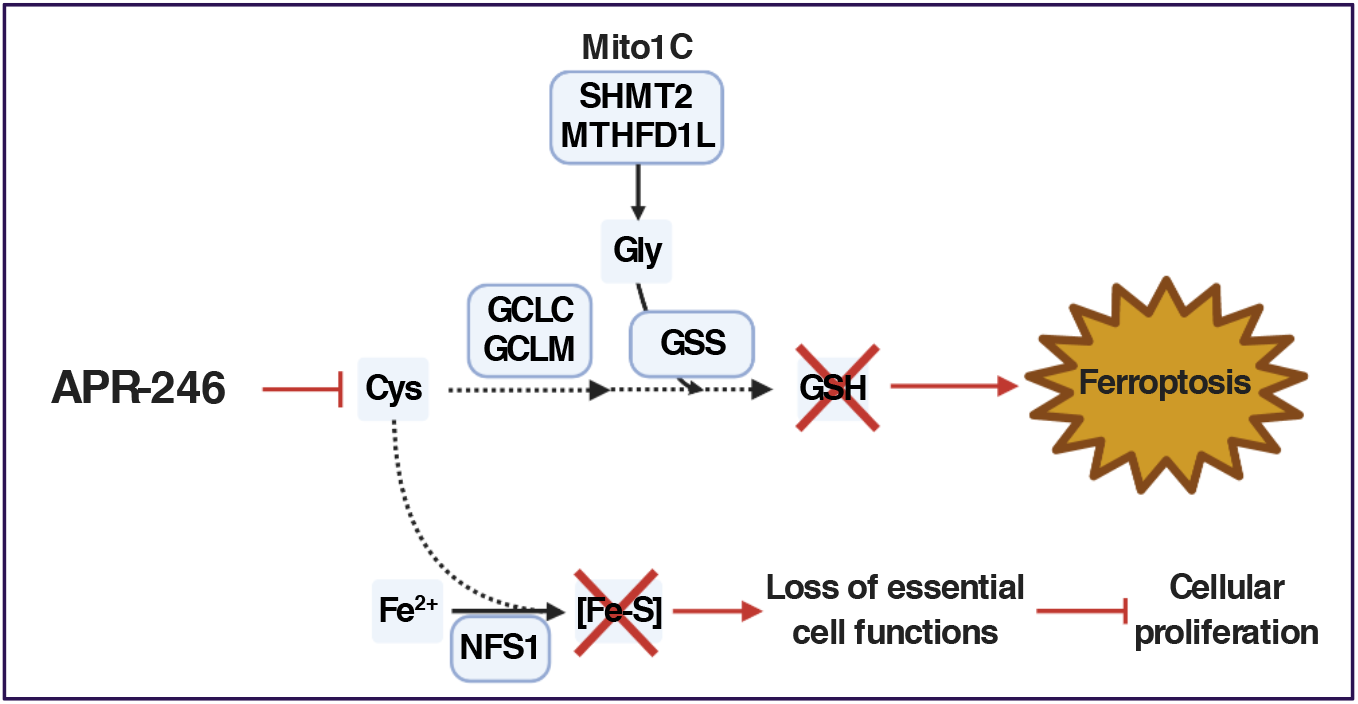

## MAIN

To distinguish the mechanism of action of APR-246, we designed and executed a set of complimentary genome-wide CRISPR perturbation screens (**Figure 1A**). We transduced OACM5.1 oesophageal cancer cells expressing Cas9-mCherry with the Brunello genome-wide CRISPR knockout (CRISPRko) library^4^ and challenged these cells with a sublethal dose of APR-246 for 8 days. In parallel, we transduced OACM5.1 cells expressing dCas9-VP64 with the Calabrese Set A genome-wide CRISPR activation (CRISPRa) library^5^ and challenged these cells with a lethal dose of APR-246 for 28 days. The CRISPRko screen aimed to identify genetic deletions that ‘drop-out’ in the APR-246 treatment cells compared to vehicle treated cells. Meanwhile, the CRISPRa screen aimed to identify gene overexpression that protects cells from APR-246. In this way, the CRISPRa screens has the advantage of findings genetic modulators of APR-246 sensitivity that may not be revealed in the CRISPRko screen as essential genes drop-out during the puromycin selection.

**Figure 1.**
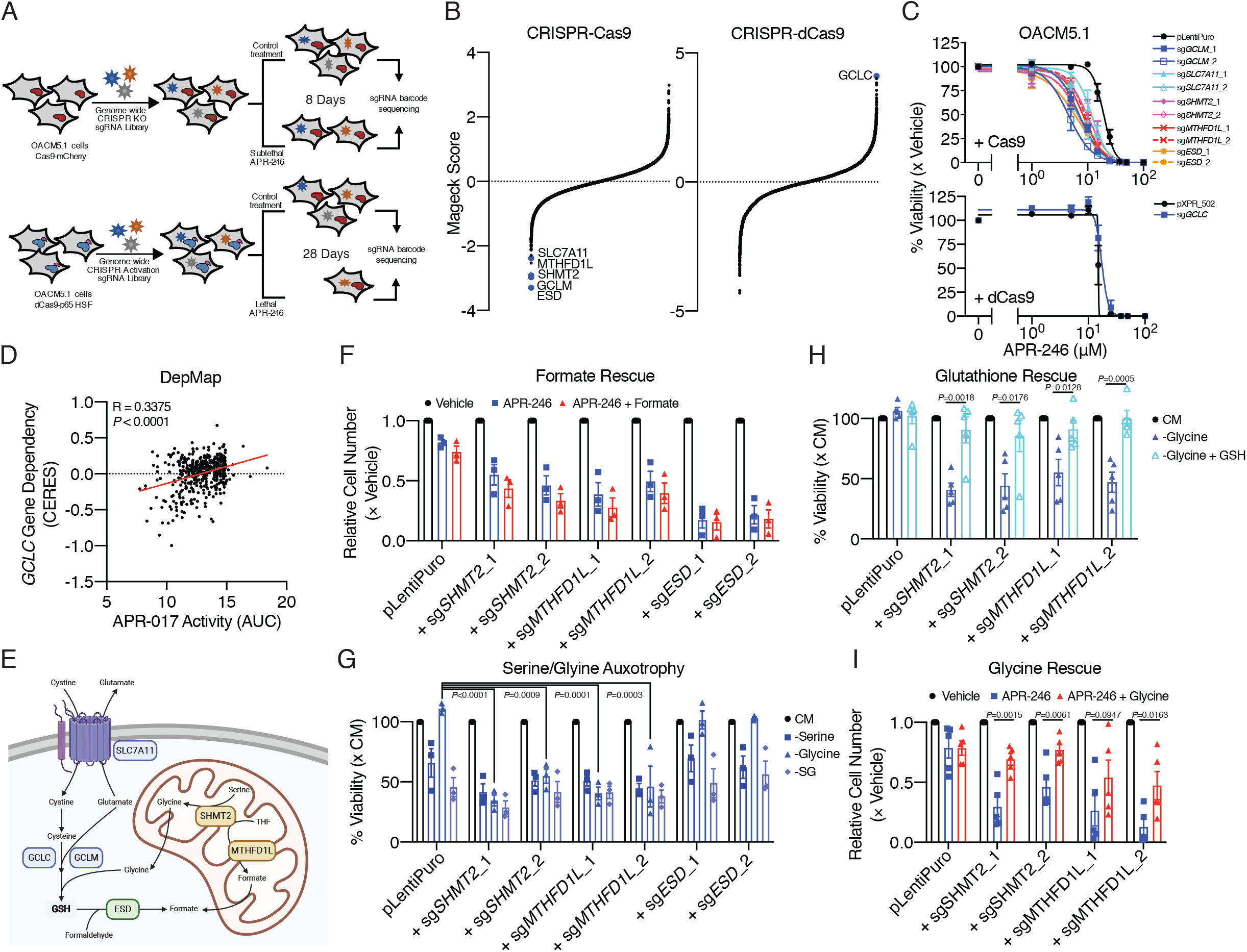
Genome-wide CRISPR screens highlight the role of *de novo* GSH synthesis and mito-1C metabolism in APR-246 mechanism of action. (A) Schematic diagram details the protocols utilised for the CRISPRko and CRISPRa screens in OACM5.1 cells. Sublethal dose = 10 μM. Lethal dose = 15 μM. (B) Mageck scores (negative indicating ‘dropout’ and positive indicating ‘enrichment’) from CRISPR screens plots in order to magnitude. (C) Cell viability following 72 h exposure with APR-246 at indicated doses to validation of hits. (D) Scatterplot correlating the expression of *GCLC* gene dependency with APR-017 (APR-246 analogue) activity (area-under the curve, AUC). (E) Schematic diagram illustrating the connections between *de novo* GSH synthesis and mito-1 metabolism. (F) Relative cell numbers of OACM5.1 cells with indicated vectors treated with 10 μM APR-246 ± 1 mM sodium formate supplementation for 4 days. (G) Cell viability following 72 h of serine, glycine or serine and glycine (SG) deprivation compared to complete media (CM). (H) Cell viability following 72 h of glycine deprivation ± 1 mM GSH-monoethyl ester (GSH-MEE). (I) Relative cell number following treatment with 10 μM APR-246 ± 1 mM glycine supplementation for 4 days. (D) Pearson’s correlation. (G) one-way ANOVA with Dunnet’s multiple comparison post-test. (H,I) unpaired two-tailed t-test. Error bars = SEM. (B) *N*=2 for CRISPRko screen, *N*=1 for CRISPRa. (C,F,G) *N*=3. (H,I) *N*=5.

Consistent with our and others prior observations that APR-246 triggers GSH depletion in cancer cells^6–9^, the CRISPRko screen identified *de novo* GSH synthesis genes *SLC7A11* and *GCLM* whilst *GCLC* was the top enriched gene in the CRISPRa screen (**Figure 1B**). Notably, two mitochondrial one-carbon (mito-1C) metabolism genes, *SHMT2* and *MTHFD1L*, and the s-formylglutathione hydrolase gene, *ESD*, involved in GSH-mediated formaldehyde detoxification were also identified^10–12^. We confirmed that two independent sgRNA guides targeting *GCLM*, *SLC7A11*, *SHMT2*, *MTHFD1L* and *ESD*, and one sgRNA guide for *GCLC* activation, modulated sensitivity to APR-246 as expected (**Figure 1C**). Consistent with this, gene dependency on *GCLC* correlated with cancer cell line (CCL) sensitivity to APR-246 analogue, APR-017 (PRIMA-1), across 456 CCLs of diverse lineage (**Figure 1D**).

We next investigated involvement of mito-1C metabolism in APR-246 sensitivity. Of note, mito-1C metabolism intersects with *de novo* GSH synthesis through SHMT2-mediated production of glycine from serine^13^, and GSH-mediated formaldehyde detoxification through MTHFD1L-mediated production of formate^10^ (**Figure 1E**). First, we attempted to rescue the increased sensitivity to APR-246 in mito-1C or *ESD* deficient cells with exogenous formate, however, formate supplementation did not alter sensitivity to APR-246 (**Figure 1F**). Previous studies have shown that mito-1C deficient cells have altered dependency on exogenous glycine due to the loss of the glycine production through SHMT2 when mito-1C is perturbed^14^. Indeed, cells expressing *SHMT2* and *MTHFD1L* sgRNAs displayed increased glycine auxotrophy whilst cells transduced with control vector or *ESD* sgRNAs did not (**Figure 1G**). Meanwhile, all cells displayed sensitivity to serine deprivation and simultaneous serine and glycine deprivation. Importantly, the glycine auxotrophy in mito-1C deficient cells could be reversed with the exogenous supply for cell permeable GSH (**Figure 1H**) - in agreement with previous studies^14^. This highlights that mito-1C deficient cells have a higher demand for exogenous glycine for *de novo* GSH synthesis. As a result, the increased sensitivity to APR-246 mito-1C deficient cells can be reversed by supplementation of additional glycine to the cell media (**Figure 1I**). Together, these data demonstrate that mito-1C metabolism and *de novo* GSH synthesis interplay to regulate the mechanism of action of APR-246.

Given the involvement of mito-1C and GSH metabolism in regulating sensitivity to ARP-246 in cancer cells, we sought to investigate whether limiting supply of serine and glycine (SG) or cystine (CySS) to cancer cells increased sensitivity to APR-246. Indeed, both SG and CySS depletion increased FLO-1 LM cells, a model of spontaneous metastatic oesophageal cacner^15^, to APR-246 in cell culture (**Figure 2A**). Restriction of dietary serine and glycine has previously been shown to reduce tumour growth in xenograft and genetically engineered mouse models^16,17^. Whilst cystine depletion *in vitro* has been widely demonstrated to induce ferroptosis^18,19^, to our knowledge, cystine restriction has not be previously shown to inhibit tumour growth. Using the FLO1 LM xenograft model in immunocompromised mice to access the effects of SG or CySS diet restriction on APR-246 efficacy, we found that SG restriction significantly enhanced the efficacy of APR-246 (**Figure 2B**). Interestingly, CySS restriction had no effect on tumour, whilst SG restriction alone had greater efficacy than APR-246 monotherapy. SG and CySS restriction in combination with APR-246 were well tolerated (**Figure 2C**) and chow consumption remained consistent across the different diet interventions (**Figure 2D**). These results demonstrate, for the first time, that SG restriction can inhibit the growth of oesophageal cancer tumours and synergies with APR-246.

**Figure 2.**
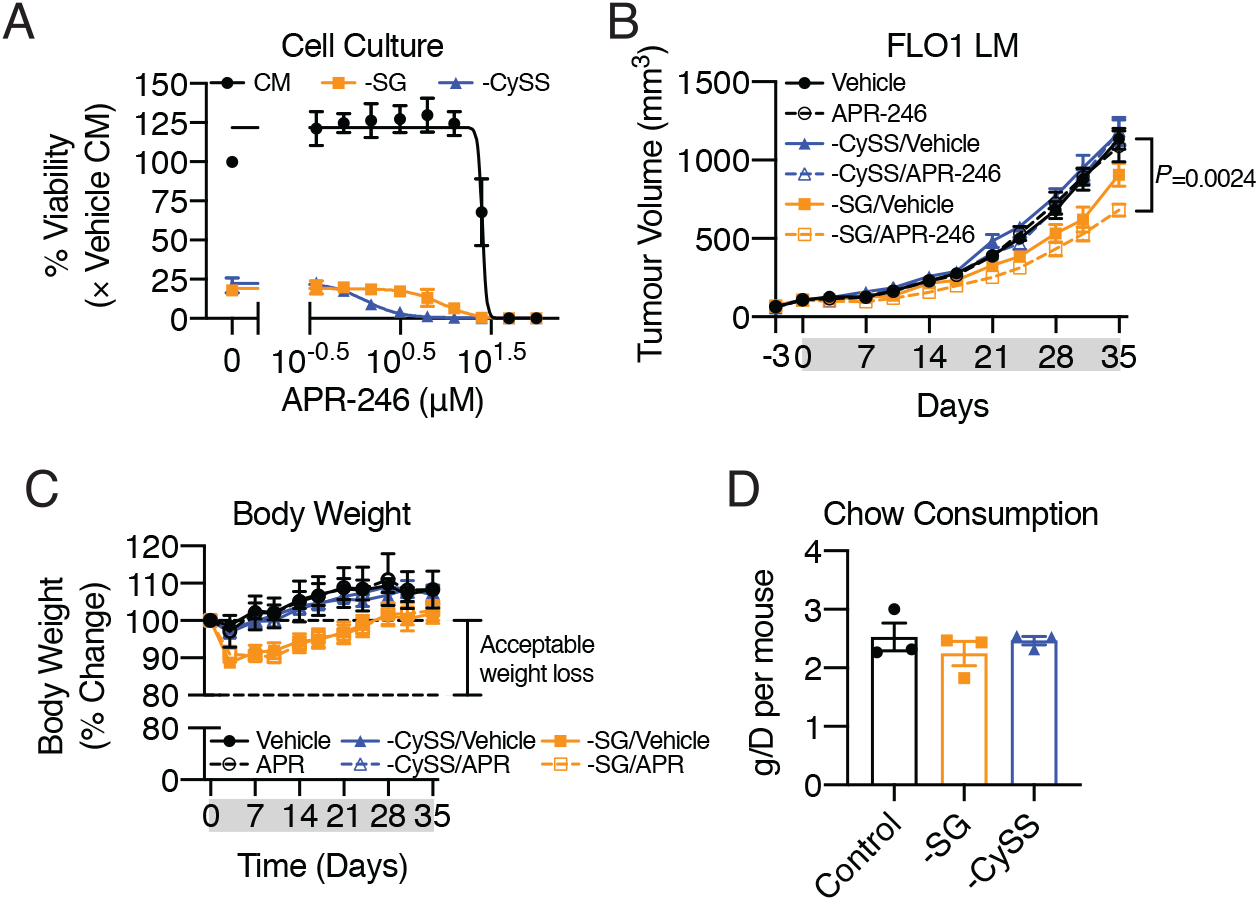
SG-free diet enhances APR-246 efficacy *in vivo*. (A) Cell viability in FLO1 LM cell following 72 h exposure to APR-246 at indicated doses in the absence of SG or cystine (CySS) compared to CM. (B) Growth curves of FLO1 LM tumours treated with APR-246 (100 mg/kg, daily) on either control, SG-free, CySS-free diets for 35 days. (C) Body weight as a percentage from baseline. Ethically acceptable weight loss is defined by Peter MacCallum Cancer Centre Animal Experimentation Ethics Committee as <20% compared to pre-treatment days. (D) Average chow consumption over 7 days per box of mice per mouse. (B) one-way ANOVS with Dunnet’s multiple comparison post-test. Error bars = SEM. (A,D) *N*=3. (B,C) *N*=8 for all groups.

Returning to our investigation the mechanism of action of APR-246, we next examined whether APR-246 induces ferroptosis. To do this, we utilised three distinct inhibitors of ferroptosis - N-acetylcysteine (NAC), ferrostain-1 (Fer-1) and ciclopirox olamine (CPX) (**Figure 3A**). Indeed, we found that APR-246-induced cell death, as measured by propidium iodine (PI) uptake, could be reversed by all the ferroptosis inhibitors, but not by the pan-caspase inhibitor, Z-VAD-FMK in both p53-null H1299 lung cancer cells and mut-p53 expressing FLO-1 oesophageal cancer cells (**Figure 3B** **- top**). Importantly however, the cellular proliferation defect induced by APR-246 was only reversed in cell co-treated with NAC (**Figure 3B** **- bottom**). This indicates that the mechanism of action of APR-246 likely diverts at the level of conjugating to cysteine to induce cell death through GSH depletion and another perturbed pathway that blocks cell proliferation. This phenomenon may also provide an explanation for why previous studies of APR-246-mediated cell death that relied on long term exposures (> 48 hr) with APR-246 and the use of cellular metabolism assays as surrogates for cellular viability failed to demonstrate rescue with ferroptosis inhibitors^6^. Furthermore, this is in keeping with recent findings that distinguish erastin, which block cystine import into cells, as maintaining cell proliferation activity in the presence of ferroptosis inhibitors^20^. Extending this, given that the proposed mechanism of action of APR-246 is to induce apoptotic cell death, we also genetically engineered the Mc38 mouse colon adenocarcinoma model to be completely deficient in the intrinsic apoptotic machinery (Bax/Bak/Bid/Casp3/Casp7 KO). We found that APR-246 was equally as efficacious in both apoptosis proficient cells and apoptosis deficient cells (**Figure 3C**). Furthermore, apoptosis deficient cells underwent cell death following exposure to APR-246 (**Figure 3D** **- left**). Conversely, staurosporine (STS) failed to induce cell death in these cells (**Figure 3D** **- right**). Together, this proves that APR-246 induces ferroptosis and does not require apoptotic machinery in order to elicit cell death.

**Figure 3.**
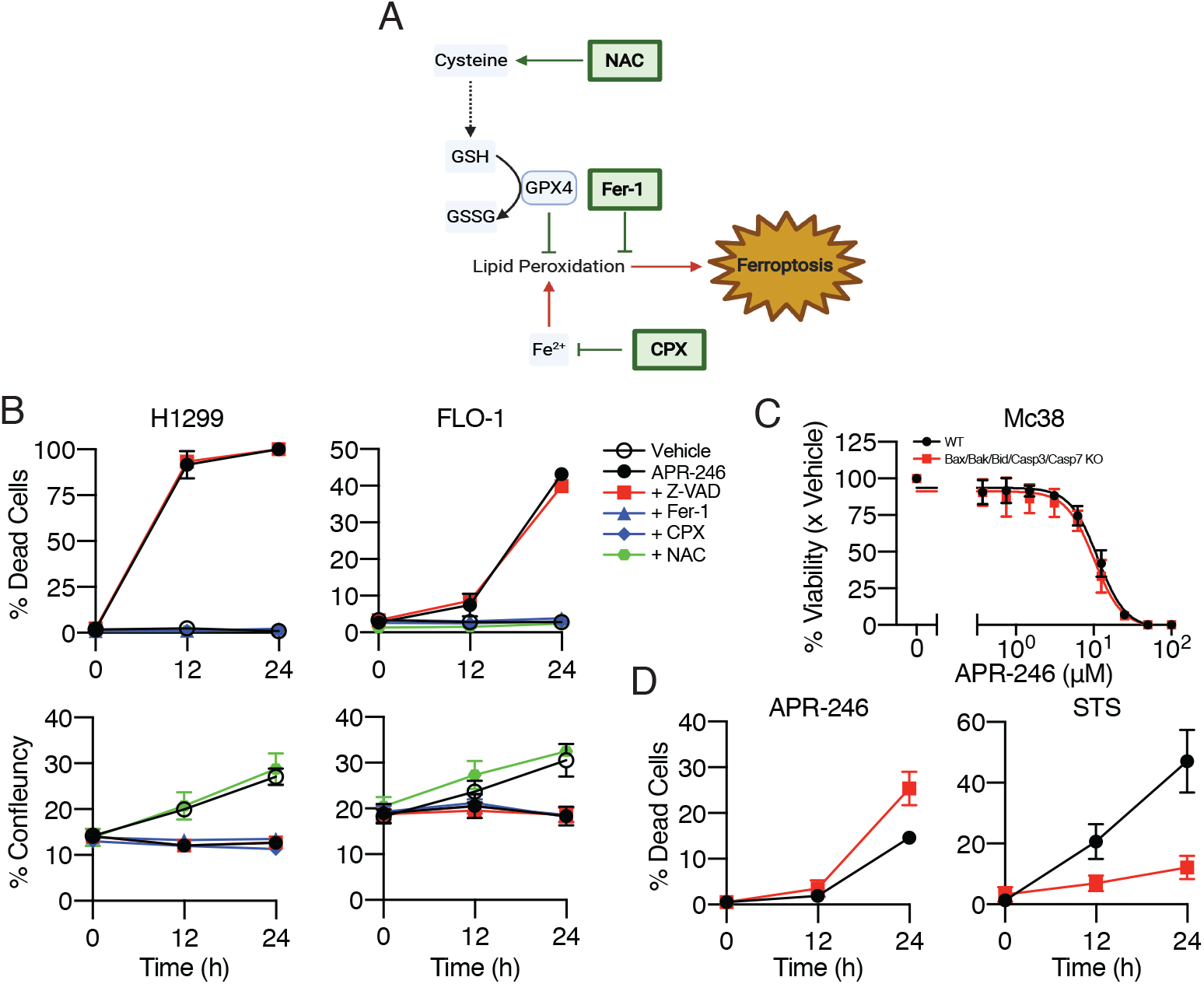
APR-246 triggers ferroptosis. (A) Schematic diagram illustrating the ferroptosis inhibitors mechanisms. (B) Percentage of dead cells (as determined by percentage of PI+ cells) and cell proliferation (as determined by cell confluency) induced by 50 μM APR-246 co-treated with 50 μM Z-VAD-FMK (pan-caspase inhibitor), 12.5 μM Ferrostain-1 (Fer-1, lipophilic antioxidant), 6.25 μM ciclopirox olamine (CPX, iron chelator) and 2.5 mM N-acetyl-cysteine (NAC, cysteine supplement) in H1299 and FLO1 cells. (C) Cell viability in Mc38 apoptosis-proficient (black, WT) and apoptosis-deficient (red, Bax/Bak/Bid/Casp3/Casp7 KO) cells. (D) Cell death induced in WT and apoptosis deficient Mc38 cells by 50 μM APR-246 or 12.5 μM Staurosporine (STS). Error bars = SEM. (B,C,D) *N*=3.

Next, we sought to explain the cell proliferation defect induced by APR-246. To this end, we performed label-free proteomics and untargeted metabolomics in OACM5.1 cells treated with APR-246 prior to the onset of ferroptosis (**Figure 4A,B**). In response to APR-246 treatment, cells upregulated mitochondrial protein ferredoxin 1 (FDX1), a critical component of mitochondrial iron-sulfur cluster biosynthesis, as well as two other protein, DNAJA3 and ABCF2, with reported roles in iron metabolism^21,22^. As expected, APR-246 treatment reduced GSH levels and simultaneously increased cystine levels likely as an adaptive response to APR-246 treatment. Meanwhile, mitochondrial metabolism altered as demonstrated by the downregulation of acetyl-CoA and NADH. As a result of these investigations, we hypothesised that APR-246 may be inhibiting mitochondrial iron-sulfur cluster biogenesis likely through binding to free cysteine and limiting the cysteine desulferase activity of NFS1 (**Figure 4C**). We first interrogated the publicly available gene essentiality data, correlating CCL dependency on *NFS1* and the activity of the 480 compounds across over 800 CCLs. Supporting our hypothesis, the top compound correlated to *NFS1* dependency was APR-246 analogue, APR-017 (**Figure 4D**). We also confirmed, in line with previous studies^23^, that MQ is able to bind directly to free cysteine (**Figure 4E**). Importantly, we shown a cell free NFS1 activity assay and in intact cells that MQ is able to inhibit the desulfurase activity of NFS1 in a dose dependent fashion (**Figure 4F,G**). Furthermore, MQ also reduced aconitase activity, a surrogate marker for iron-sulfur stability^24^ (**Figure 4H**). Together, these results demonstrate that APR-246 inhibits iron-sulfur cluster biogenesis through cysteine conjugation - providing a novel explanation for the cell proliferation defect induced by APR-246 in the presence of ferroptosis inhibitors.

**Figure 4.**
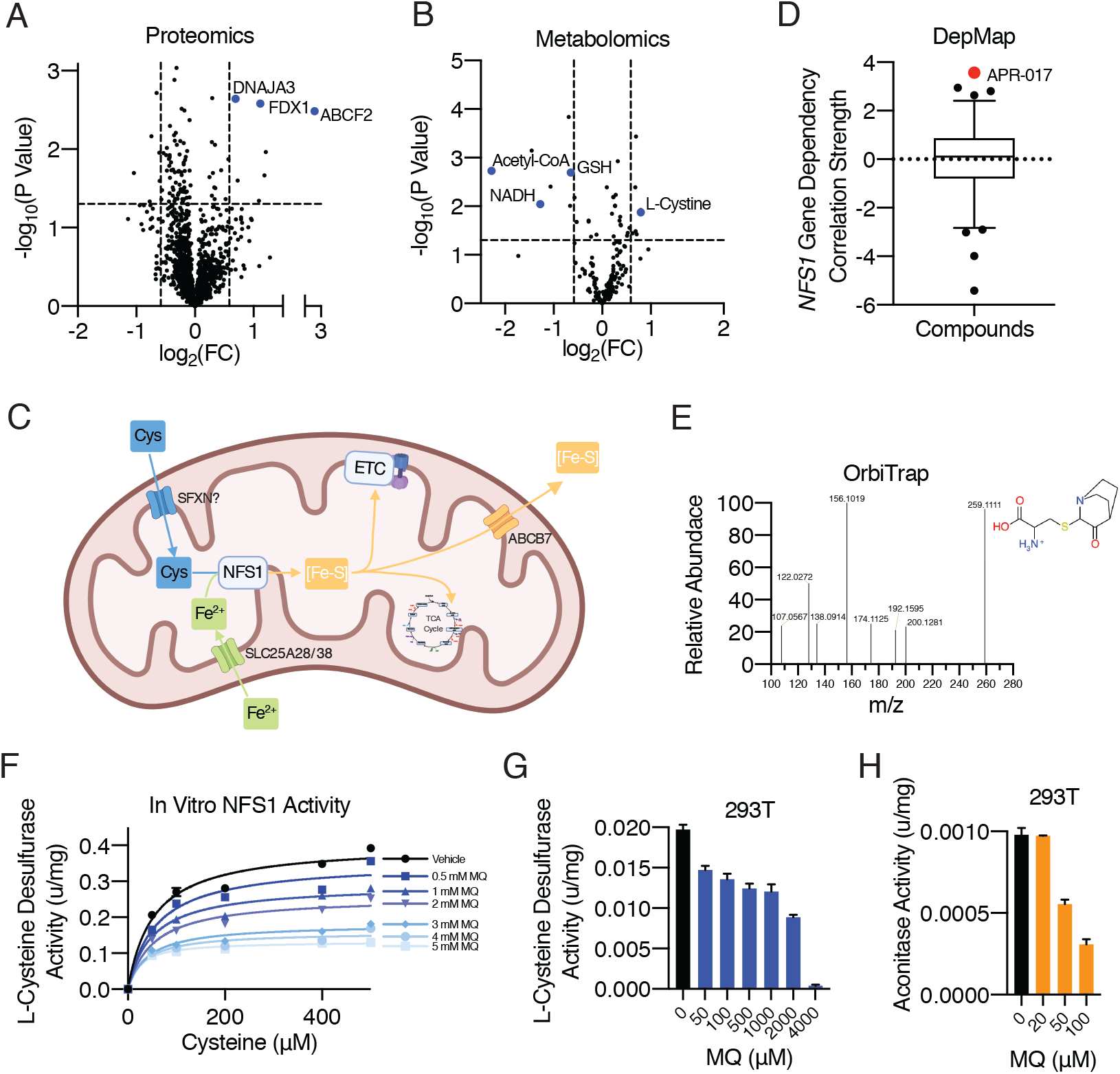
APR-246 inhibits iron-sulfur cluster biogenesis. (A) Changes in proteins determined by label-free quantitative proteomics and (B) untargeted LC-MS metabolomics following 50 μM APR-246 for 12 h compared to vehicle. (C) Schematic diagram illustrating mitochondrial iron-sulfur cluster biogenesis and its role in mitochondrial metabolism. (D) Box-and-whisker plot (1st-99th percentile) of Fischer’s transformed z-scored Pearson correlation strength of *NFS1* gene dependency and the 481 Cancer Therapeutic Response Portal v2 (CTRPv2) compounds. Red dot indicates APR-017 (APR-246 analogue) is the top positive correlated compound to *NFS1* gene dependency. (E) Mass spectrometry pattern of 10 μM pre-heated APR-246 mixed with 10 μM cysteine in H2O showing MQ conjugated to cysteine (MQ-Cys, m/z = 259.111 [m + H]). (F) Cell-free and (G) in-cell (in HEK-239T cells) L-cysteine desulfurase assay measuring NFS1 activity at indicated doses of MQ following 1 h incubation. (H) Aconitase activity assay in HEK-239T cells following 1 h incubation with MQ at indicated doses. Error bars = SEM. (A) *N*=4. (B) *N*=6. (F,G,H) *N*=3.

In sum, we show that APR-246 induces ferroptosis and inhibits iron-sulfur cluster biogenesis through cysteine conjugation. This has broad implications for the clinical utility of APR-246. This proposed mechanism of action highlights that APR-246 should be expanded to beyond mutant-p53 cancers, which we have illustrated in previous studies^25^. In the context of *TP53*-mutated MDS, APR-246 is currently only indicated in ~7% of all MDS cases^26^. Dysregulated iron metabolism is a hallmark of a subset of MDS with ringed sideroblasts, characterised by aberrant iron accumulation in erythroblast mitochondria^27^. The aberrant iron accumulation in MDS with ringed sideroblasts likely increases erythroblasts sensitivity to ferroptosis. Importantly, the most frequent genomic alteration associated with MDS with ringed sideroblasts are *SF3B1* mutations, however these are mutually exclusive with *TP53* mutations^28^. Therefore, our work reveals that the clinical utility of APR-246 could be potentially expanded to include MDS cases with aberrant iron accumulation in order to leverage its capacity to induce ferroptosis.

## METHODS SUMMARY

CRISPR screens performed in human OACM5.1 cells. Validation was performed on polyclonal OACM5.1 cells following selection with puromycin. Formate and glycine rescue experiments were performed in conditions matching the CRISPR screening procedure scaled down to T25 flask. Serine/Glycine auxotrophy experiments were perform in 96-wells plates utilising dialyzed FBS and media reconstituted without serine and glycine. Cancer dependency data and APR-017 cancer cell line sensitivity data was obtained from the DepMap web-portal (www.depmap.org) and correlation analyses were performed on R studio using the cor.test function. For diet intervention studies, ~6-week-old female NSG mice were sub-cutaenously injected with 5×10^6^ FLO1 LM cells and randomised to SG-deplete, CySS-deplete or control chow (AIN93G Rodent Diet) and dosed with 0.9% saline or 100 mg/kg APR-246 (Aprea) once tumours reach 100mm^3^. For ferroptosis rescue experiments, cell death and proliferation were quantified utilising IncuCyte FLR (Essen BioSciences). Cell viability was determined by alamarBlue assay (Life Technologies) using Cytation 3 (BioTek). Mc38 apoptosis-deficient cells were generated using CRISPR-Cas9 technology. Label-free proteomics (LC-MS/MS) was performed at the Mass Spectrometry and Proteomics Facility at Bio21 Institute and untargeted metabolomics (LC-MS) in collaboration with Metabolomics Australia. NFS1 cell-free and in cell assays and aconitase assays were performed as previously described^29^.

